# Heroes and villains: opposing narrative roles engage neural synchronization in the lateral inferior frontal gyrus

**DOI:** 10.1101/2023.08.24.554721

**Authors:** Hayoung Ryu, M. Justin Kim

## Abstract

Neuroscientific studies have highlighted the role of the default mode network (DMN) in processing narrative information. Here, we examined whether the neural synchronization of the DMN tracked the appearances of characters with different narrative roles (i.e., protagonists versus antagonists) when viewing highly engaging, socially rich audiovisual narratives. Using inter-subject correlation analysis on two independent, publicly available movie-watching functional magnetic resonance imaging datasets (*Sherlock* and *The Grand Budapest Hotel*), we computed whole-brain neural synchronization during the appearance of the protagonists and antagonists. Results showed that the inferior frontal gyrus (IFG) and orbitofrontal cortex (OFC), which are components of the DMN, had higher ISC values during the appearance of the protagonists compared to the antagonists. Importantly, these findings were commonly observed in both datasets. We discuss the present results in the context of information integration and emotional empathy, which are relevant functions known to be supported by the DMN. Our study presents generalizable evidence that regions within the DMN – particularly the IFG and OFC – show distinctive synchronization patterns due to differences in narrative roles.

## Introduction

We commonly experience and immerse in numerous narratives as a natural part of our daily routine by reading, watching, and listening to stories. Just as in real-life human interactions, readers and audiences actively infer the actions, goals, and intentions of characters to understand them. People attempt to understand the constructed reality of a narrative and share the feelings with the characters, at least in part, through experience-taking, a process in which the perceivers replace part of their identity with the mindset of the characters (Kaufman & Libby, 2012). For example, people who tend to identify with the protagonists of a narrative are more susceptible to vicarious emotional experience such as stress (Tannenbaum & Gaer, 1965). This experience shares similar qualities with empathy – the sharing of the feelings of another person (Decety & Jackson, 2004). Indeed, there is a growing body of research that links narrative engagement to social cognition (Mar, 2018; Maslej et al., 2017; DilllJShackleford et al., 2016; Tamir et al., 2016; Mar & Oatley, 2008). These studies imply that narrative engagement may have real-world impact via long-lasting positive socioemotional outcomes. For example, exposure to fiction improves the capacity for empathy and social inference (Kidd & Castano, 2013; Black & Barnes, 2015), and can even affect functional brain connectivity in both short- and long-term (Berns et al., 2013). As such, narrative engagement is beyond a form of entertainment; it shapes a person’s experience with characters, and in turn significantly influences the manner in which they navigate their social worlds.

Given that narrative engagement is a popular multidisciplinary research topic, it is noteworthy that there are relatively few neuroimaging studies that examines the neural basis of narrative-driven character perception. One recent functional magnetic resonance imaging (fMRI) study found that the activity of the default mode network (DMN) distinguishes popular fictional characters based on their narrative roles (Ron et al., 2022). Classification analysis revealed that brain regions overlapping with the DMN showed better performance than the occipital face area, indicating that social categorization based on narrative roles engages the DMN. Another recent fMRI work by Ohad and Yeshurun (2023) examined individual differences in neural synchronization as a function of engagement to the narrative and characters. According to this study, the level of engagement was related to synchronization within the DMN that included the dorsomedial prefrontal cortex (dmPFC), temporoparietal junction (TPJ) and right temporal pole as key nodes. Of relevance to the present study, Ohad and Yeshurun (2023) observed a dissociation in neural synchronization patterns during the appearance of positively and negatively engaging characters. Both fMRI studies, using different measurements and analytical strategies, offer converging evidence that highlight the DMN in distinguishing different characters based on their narrative roles.

The importance and contributions of these fMRI studies notwithstanding, there are some notable limitations that could be improved upon. For instance, Ron and colleagues (2022) used static images of the fictional characters from different movies to elicit brain activity, precluding potential effects of ongoing perception of actions, movements, and dynamic facial expressions, as well as any in-narrative contexts. As suggested by previous studies, using naturalistic stimuli such as movies, music, and audiobooks improves ecological validity compared to traditional static visual stimuli in neuroimaging studies (Nastase et al., 2019, 2020; van Atteveldt et al., 2018, Klin et al., 2002). Through the use of such naturalistic stimuli, it is especially important and useful to examine the complex nature of socioemotional processing by capturing how our brains function in the wild (Saarimäki, 2021; Tikka et al., 2023).

In the present study, we aimed to investigate the involvement of brain regions, the DMN in particular, as participants watched complex and socially rich audiovisual narratives using two independent, publicly available movie-watching fMRI datasets. Under the assumption that people gain experience through the lens of characters as a story unfolds (Kaufman & Libby, 2012), there would be measurable differences in how they process each narrative role depending on how relatable the character is, or how much information they have about the character. We reasoned that the characters with the most prominent and distinctive narrative roles – protagonists and antagonists – would offer a useful window into the neural basis of such processes. Using fMRI, we adopted the inter-subject correlation (ISC) method to test the neural synchronization among individuals while they experienced the same time-locked stimuli (Hasson et al., 2004). The basic premise of the ISC analysis is that, as the subjects watch the same movie, the blood-oxygen-level-dependent (BOLD) signal in brain regions that putatively process the stimuli will systematically synchronize across subjects. On the other hand, neural activity in the brain regions that do not respond to the stimuli will be random, leading to little to no synchronization. Therefore, ISC values can be interpreted as how consistent the region is involved in processing the stimuli (Nastase et al., 2019; Nguyen et al., 2019; Grall et al., 2021). In the present study, we posit that there will be different neural synchronization pattern, indexed by ISC, based on the most prominent and contrasting narrative roles – protagonists and antagonists. We hypothesized that the DMN will be involved during the processing of different narrative roles, characterized by separable neural synchronization patterns during the appearance of protagonists and antagonists. To increase the generalizability of our findings, we sought to identify and report brain regions that were reliably observed across two independent datasets.

### Methods Datasets

We utilized two open fMRI datasets that leveraged naturalistic stimuli with a narrative focus. The first dataset comprises fMRI scans of 16 subjects (demographic information was only available for the initial 22 participants of the original study; mean age = 20.8, ages 18-26, 12 males, 10 females) watching the initial 50 minutes of the 2010 BBC TV series *Sherlock* (Chen et al., 2018). The second dataset contains fMRI scans of 25 subjects (mean age = 27.5, ages 22-31, 13 females, 12 males) watching the second half of the 2014 film *The Grand Budapest Hotel* (di Oleggio Castello et al., 2020). Both datasets are available to download at OpenNeuro (*Sherlock*: https://openneuro.org/datasets/ds001132; *The Grand Budapest Hotel*: https://openneuro.org/datasets/ds003017). We chose these two datasets as both *Sherlock* and *The Grand Budapest Hotel* contain socially rich contents with an engaging narrative structure that centers around its main characters. Our focus was on the main protagonist and antagonist in each story, which corresponds to Sherlock and Mycroft (*Sherlock*), and Gustave and Dmitri (*The Grand Budapest Hotel*).

### Preprocessing

We aimed to keep the preprocessing steps as consistent as possible across the two datasets. For *Sherlock*, we used the preprocessed fMRI data that were published along with the rest of the *Sherlock* dataset (Chen et al. 2018). In brief, Chen et al. (2018) performed realignment to MNI152 space and spatial normalization using fMRIPrep (Esteban et al., 2019). Spatial smoothing was conducted using a 6-mm full-width at half-maximum kernel, and denoising was performed using a general linear model (GLM). Specifically, denoising involved removing global and multivariate spikes, average CSF activity, linear and quadratic trends, as well as 24 motion covariates. Among the initial 22 participants, six of them were discarded due to excessive head motion (*n* = 2 participants), short recall data (*n* = 2), falling asleep (*n* = 1), and missing data (*n* = 1).

Since *The Grand Budapest Hotel* dataset (di Oleggio Castello et al., 2020) only provided raw data, we performed preprocessing using an equivalent pipeline as the *Sherlock* dataset via fMRIPrep. The preprocessing steps included realignment, spatial normalization, smoothing, and denoising as described above. Framewise displacement (FD) was calculated for each functional run, and two subjects (sub-sid000034, sub-sid000055) were excluded due to excessive mean FD (Power et al., 2014). The final sample size consisted of 39 (16 from *Sherlock* and 23 from *The Grand Budapest Hotel* datasets) participants. For both datasets, we used the Schaefer-Yeo atlas (Schaefer et al., 2018) to parcellate the fMRI data into 100 regions, from which mean BOLD timeseries were extracted.

### Annotation and Scene Selection

Rather than using the entire BOLD timeseries, we focused on the inter-subject synchronization of brain activity that was time-locked to each character’s appearance (Figure 1). In *Sherlock* and *The Grand Budapest Hotel*, the protagonists are Sherlock and Gustave, whereas the antagonists are Mycroft and Dmitri, respectively. It is worth noting that although Mycroft is not the primary antagonist throughout the entire episode, he is considered as such due to his actions of threatening another main character, John Watson, while self-identifying as an ‘enemy’ of Sherlock. This intentionally misleading portrayal confuses the audience, as the true criminal is not revealed within the first 50 minutes of the movie. Thus, the narrative role of Mycroft, at least in the first 50 minutes of the first episode, is consistent with that of an antagonist.

**Figure 1.**
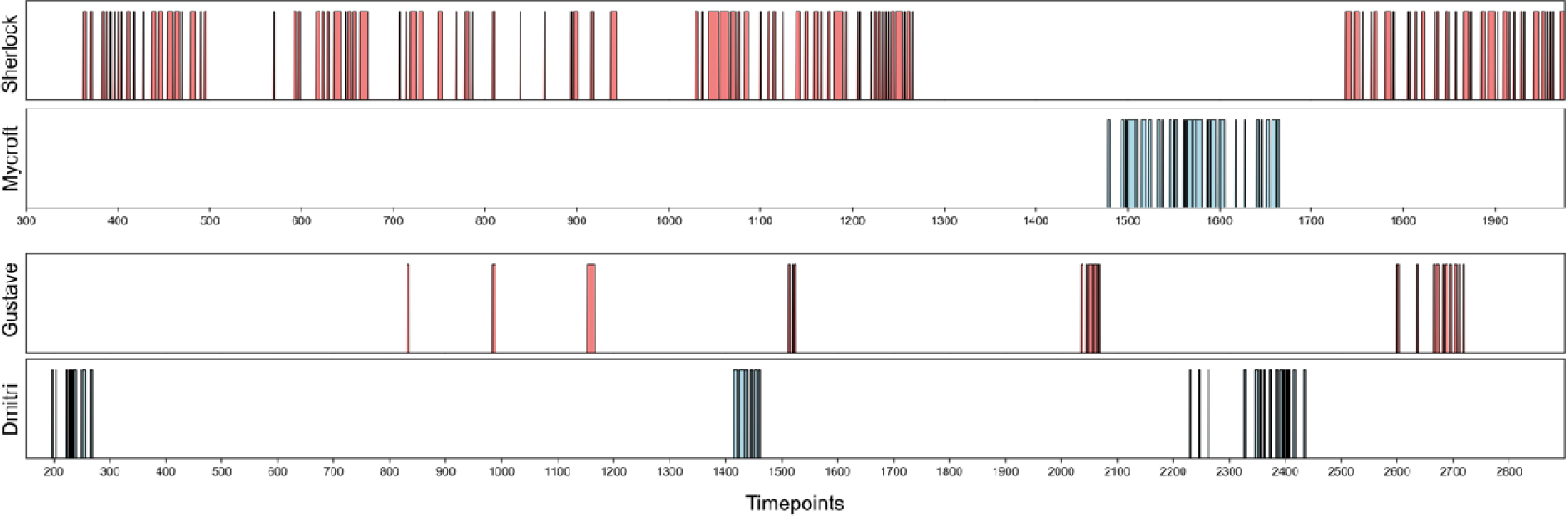
Timepoints during which the protagonists (Sherlock, Gustave) and antagonists (Mycroft, Dmitri) appeared on screen in their respective movies (first episode of *Sherlock,* second half of the film *The Grand Budapest Hotel*).

We annotated the TRs corresponding to the appearance of each character’s face alone on the screen or when the character was alongside other characters, with the camera was focused on the specific target character (e.g., camera following the protagonist/antagonist as they moved). To ensure accuracy, we excluded scenes where the protagonists and antagonists interacted in the same scene. Considering the hemodynamic delay of 3-5 seconds, we selected TRs that were 4.5 seconds (3 TRs) after the characters appeared on screen for *Sherlock* and 5 seconds (5 TRs) for *The Grand Budapest Hotel* (Hasson et al., 2004, Chang et al., 2021). The annotated time length for each character is 690 s for Sherlock, 123 s for Mycroft, 89 s for Gustave, and 92 s for Dmitri. We note that since Gustave mostly appeared with other characters, especially a sidekick character named Zero Moustafa, Gustave had fewer annotated timepoints compared to Sherlock. On the other hand, the antagonists, Mycroft and Dmitri, had similar length of appearance in their respective movies.

### ISC Calculation and Statistical Evaluation

In the present analysis, we adopted inter-subject correlation (ISC) as an index of inter-subject synchronization of brain activity. There are two major methods used to calculate ISC: pairwise or leave-one-out approach (Nastase et al., 2019). The pairwise approach involved calculating *n*(*n*-1)/2 correlation values for each voxel or parcel. In contrast, the leave-one-out approach computes ISC maps for each subject by correlating their BOLD timeseries with the average timeseries of the remaining subjects. Here, we chose the pairwise approach to account for individual variability in BOLD timeseries data, as the leave-one-out approach averages the activity time courses, potentially overlooking such variability. The synchronization of the parcels is summarized by the mean or median of the pairwise correlation values. This process is repeated for all protagonists and antagonists.

The next step is to evaluate the significance of the calculated ISC values. We adopted subject-wise bootstrapping, a non-parametric method that is often used when assumption of independence is violated during pairwise correlation approach. Subject-wise bootstrapping involves randomly sampling *N* subjects with replacement multiple times to create a bootstrap distribution. In this paper, we performed statistical evaluations using bootstrapping with 5,000 samples. We used the median to represent ISC values for each parcel since bootstrapping samples with replacement, meaning it could sample the same participant, may lead to absurdly high correlations. Therefore, the median is the preferred choice when using subject-wise bootstrapping (Chen et al., 2016; Nastase et al., 2019). As we are testing across 100 parcels, we applied Bonferroni correction (*p* < .0005) for thresholding.

### ISC Comparison between Protagonists versus Antagonists

Fisher’s *z* transformation was performed prior to statistically comparing ISCs for each parcel. As the characters had different lengths of TRs, we randomly sampled TRs from characters with more TRs (Sherlock and Dmitri) to match the number of TRs of characters with less TRs (Mycroft and Gustave). We then employed paired *t*-tests to compare ISCs across conditions (protagonists vs. antagonists) per parcel (Figure 2). This allowed us to determine which condition (i.e., protagonist or antagonist) produced greater synchronization in each parcel. To ensure that the results of the paired *t*-tests were not driven by TR selection, we repeated the process of randomly sampling TRs, ISC calculation, and paired *t*-tests over 1,000 iterations. We note that this procedure allowed us to focus on the neural responses to the perception of each character, regardless of narrative development (e.g., sampled TRs may be a collection of frames from different parts of the story, rather than a single coherent scene) – which may be a topic of scientific inquiry on its own right, but beyond the scope of the present study. If ISCs between characters differed significantly in more than 950 iterations (i.e., consistently different in > 95% of observations), we considered that parcel to synchronize differently between protagonists versus antagonists. Follow-up ISC analysis was conducted using Neurosynth parcellation (de la Vega et al., 2016) to consider involvement of amygdala, which was not covered in Schaefer-Yeo cortical parcellation. The amygdala was examined because it is often related to emotional processing (Baxter & Croxson, 2012; Adolphs et al., 1995; Mattavelli et al., 2014). Examining the amygdala was relevant as we assumed that emotional responses in some capacity is likely to accompany these characters, since the conflict between protagonists and antagonists is one of the major driving forces of any narrative. This procedure was performed separately for the *Sherlock* and *The Grand Budapest Hotel* datasets.

**Figure 2.**
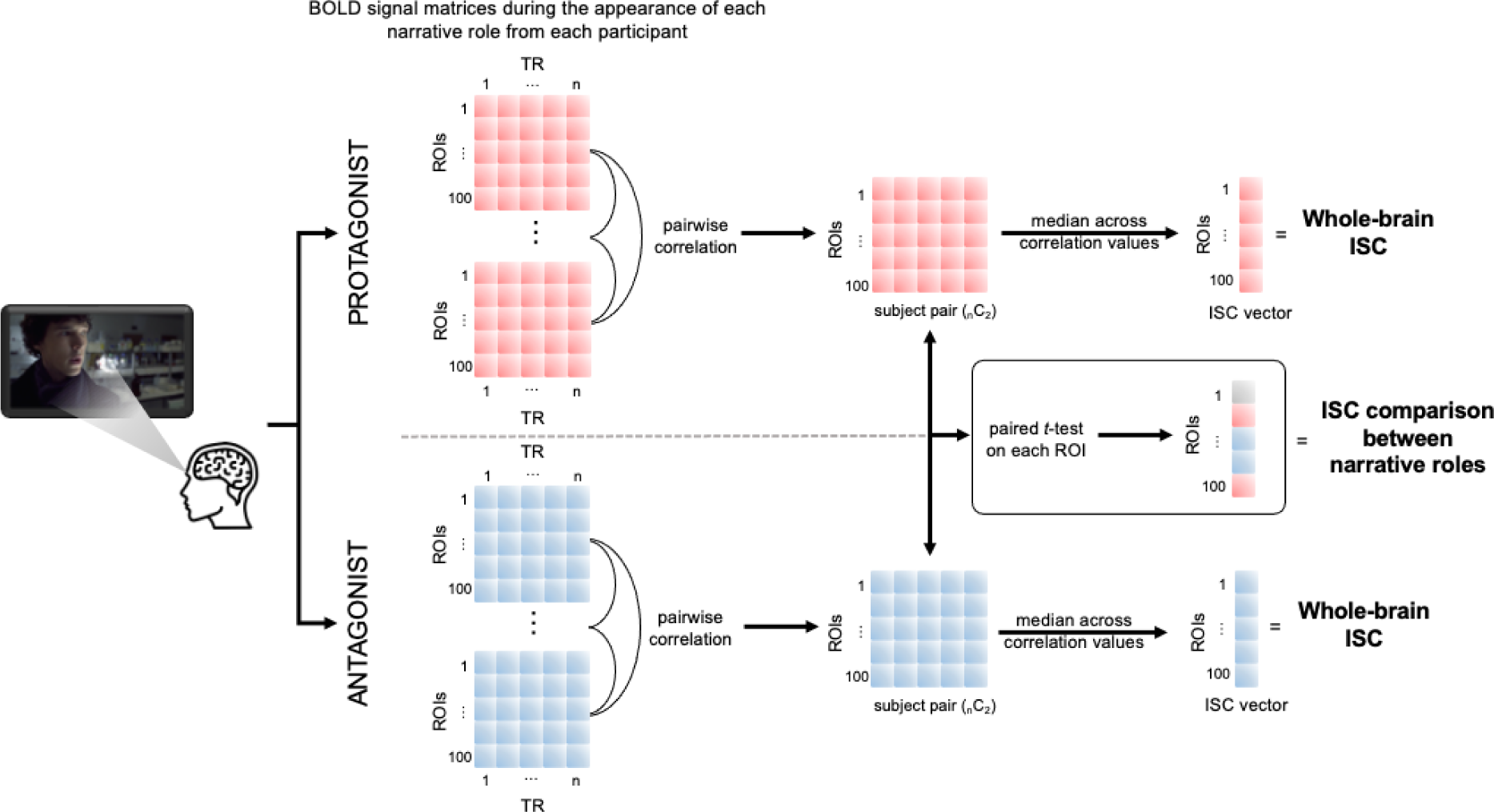
To calculate whole-brain ISC values, we computed pairwise Pearson correlations of BOLD time course data for the 100 ROIs based on the annotated TRs of each character. Median pairwise ISC values in each ROI became the final ISC value of the ROI. To compare the ISC values between narrative roles, TRs of the characters with longer screen time (i.e., Sherlock and Dmitri) were randomly sampled within the annotated TRs to match the number of TRs of the characters with shorter screen time (i.e., Mycroft and Gustave). After computing pairwise correlations of BOLD time course data, paired *t*-tests between the narrative roles were performed on an ROI-by-ROI basis. Random sampling of the TRs, ISC calculation, and paired *t*-tests were repeated for 1,000 iterations to ensure that the results were consistent regardless of TR selection. We considered the ROI to be differentially synchronized if the result was observed in 95% or more iterations.

## Results

### Whole-brain ISC

We performed whole-brain inter-subject correlation (ISC) analysis, as described by Hasson and colleagues (2004), to identify regions that were engaged during the appearances of antagonists and protagonists in both the *Sherlock* and *The Grand Budapest Hotel* datasets. As noted above, in the first episode of *Sherlock*, Sherlock and Mycroft effectively served as the protagonist and the antagonist, respectively. Whole-brain ISC analysis revealed a total of 93 regions that demonstrated statistically significant synchronization during the appearance of Sherlock (Supplementary Table S1). When categorized into canonical functional networks (Schaefer et al., 2018), 17 regions were assigned to the visual network, 9 to the somatomotor network, 15 to the dorsal attention network, 12 to the ventral attention network, 5 to the limbic network, 12 to the control network, and 23 to the default network (Figure 3). On the other hand, 57 regions were shown to synchronize during the appearance of Mycroft (Supplementary Table S2), comprising 16 regions from the visual network, 6 from the somatomotor network, 10 from the dorsal attention network, 2 from the ventral attention network, 1 from the limbic network, 7 from the control network, and 15 from the default network (Figure 3). In the additional follow-up analysis, we found that the ISC values of the amygdala during the appearance of Sherlock was not significant (ISC = 0.073, *ns*). Amygdala ISC was significant for Mycroft only at an uncorrected threshold (ISC = 0.078, *p* < .05).

**Figure 3.**
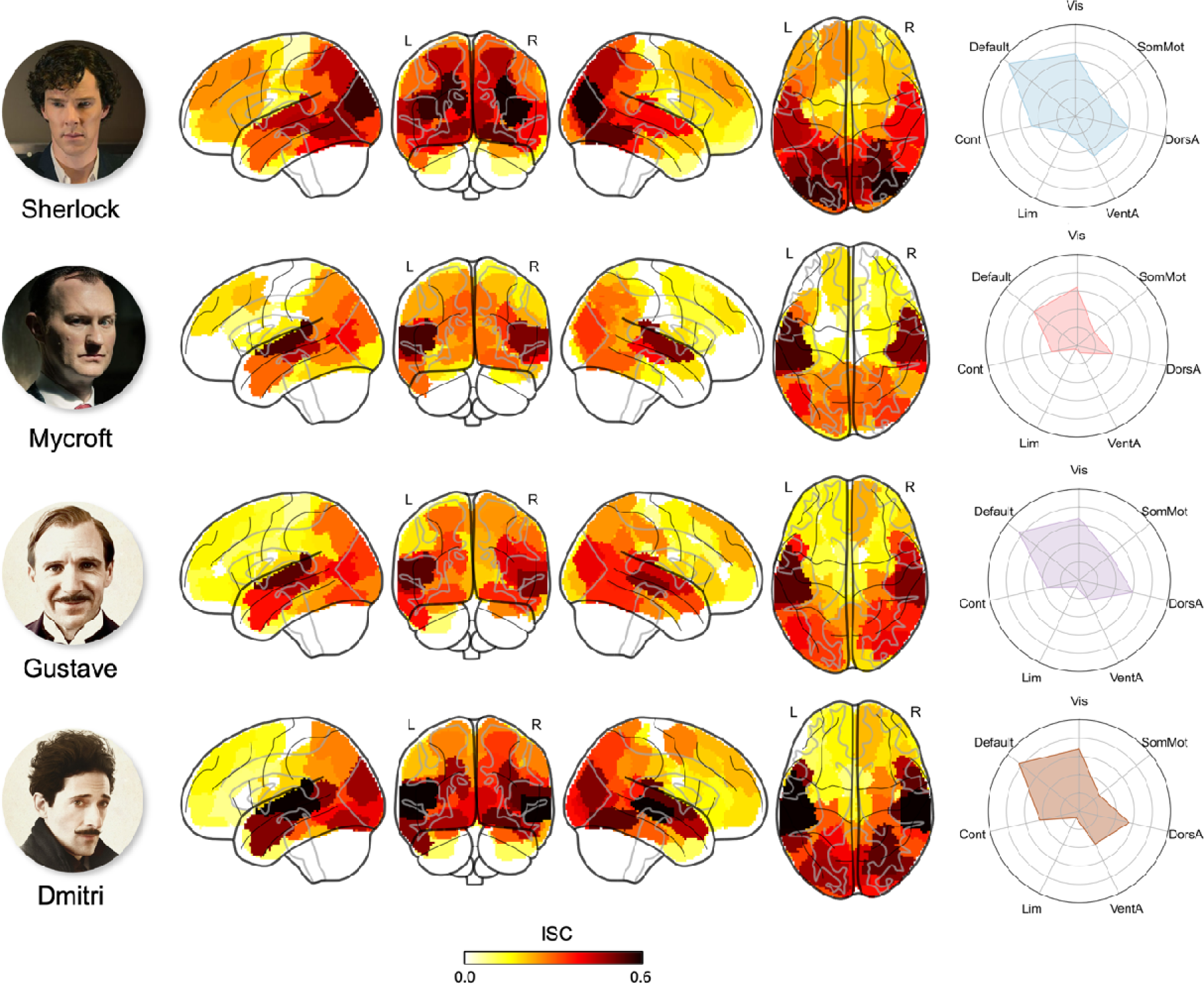
Results of the whole-brain ISC analysis during each of the character’s appearance (Bonferroni-corrected *p* < .0005). Brain regions with significant ISC values were categorized into Yeo 7 networks. SomMot = Somatomotor, DorsA = Dorsal attention, VentA = Ventral attention, Lim = Limbic, Cont = Control.

In *The Grand Budapest Hotel* dataset, whole-brain ISC analysis showed that 81 regions synchronized during the appearance of Gustave (Supplementary Table S3). These were assigned to the canonical functional networks as follows: 17 regions to the visual network, 11 to the somatomotor network, 15 to the dorsal attention network, 12 to the ventral attention network, 5 to the limbic network, 12 to the control network, and 23 to the default network (Figure 3). Conversely, during the appearance of Dmitri, 82 regions showed significant synchronization (Supplementary Table S4). Among them, 17 regions were assigned to the visual network, 7 to the somatomotor network, 14 to the dorsal attention network, 10 to the ventral attention network, 2 to the limbic network, 11 to the control network, and 21 to the default network (Figure 3). Follow-up analysis showed that both amygdala ISC values during the appearance of Gustave and Dmitri were not significant (Gustave ISC = 0.068, *ns*; Dmitri ISC = 0.044, *ns*).

### Generalizable ISC Differences Between Protagonists versus Antagonists

To determine regions with significantly different ISC values, we conducted 1,000 iterations of paired *t*-tests for each parcel with the length of appearance controlled, given that each character from each dataset appeared on screen for different durations. If the results consistently showed significant differences between characters (Sherlock vs. Mycroft and Gustave vs. Dmitri) in over 950 iterations (i.e., > 95% observations), that parcel was considered to exhibit distinctive synchronization based on narrative roles (protagonist vs. antagonist). Across both datasets, we observed four parcels that consistently displayed different synchronization patterns between characters with opposite narrative roles (Figure 4; see Supplementary Tables S5 and S6 for a complete list). Of these four, parcels 42 and 43 were part of the default network and exhibited higher ISC during the appearance of the protagonists compared to the antagonists (parcel 42: Sherlock ISC = 0.123, *p* = .0002; Mycroft ISC = 0.010, *ns*; Gustave ISC = 0.163, *p* = .0002; Dmitri ISC = 0.061, *ns*; parcel 43: Sherlock ISC = 0.217, *p* = .0002; Mycroft ISC = 0.072, *ns*; Gustave ISC = 0.145, *p* = .0002; Dmitri ISC = 0.036, *ns*)(Figure 5). Conversely, the other two parcels, parcels 10 and 59, were part of the somatomotor network and showed higher ISC during the appearance of the antagonists compared to the protagonists (parcel 10: Sherlock ISC = 0.439, *p* = .0002; Mycroft ISC = 0.543, *p* = .0002; Gustave ISC = 0.527, *p* = .0002; Dmitri ISC = 0.620, *p* = .0002; parcel 59: Sherlock ISC = 0.383, *p* = .0002; Mycroft ISC = 0.494, *p* = .0002; Gustave ISC = 0.426, *p* = .0002; Dmitri ISC = 0.522, *p* = .0002)(Figure 5). The results of follow-up analysis in both *Sherlock* and *The Grand Budapest Hotel* showed that the amygdala ISC values between the protagonists and antagonists were not significantly different.

**Figure 4.**
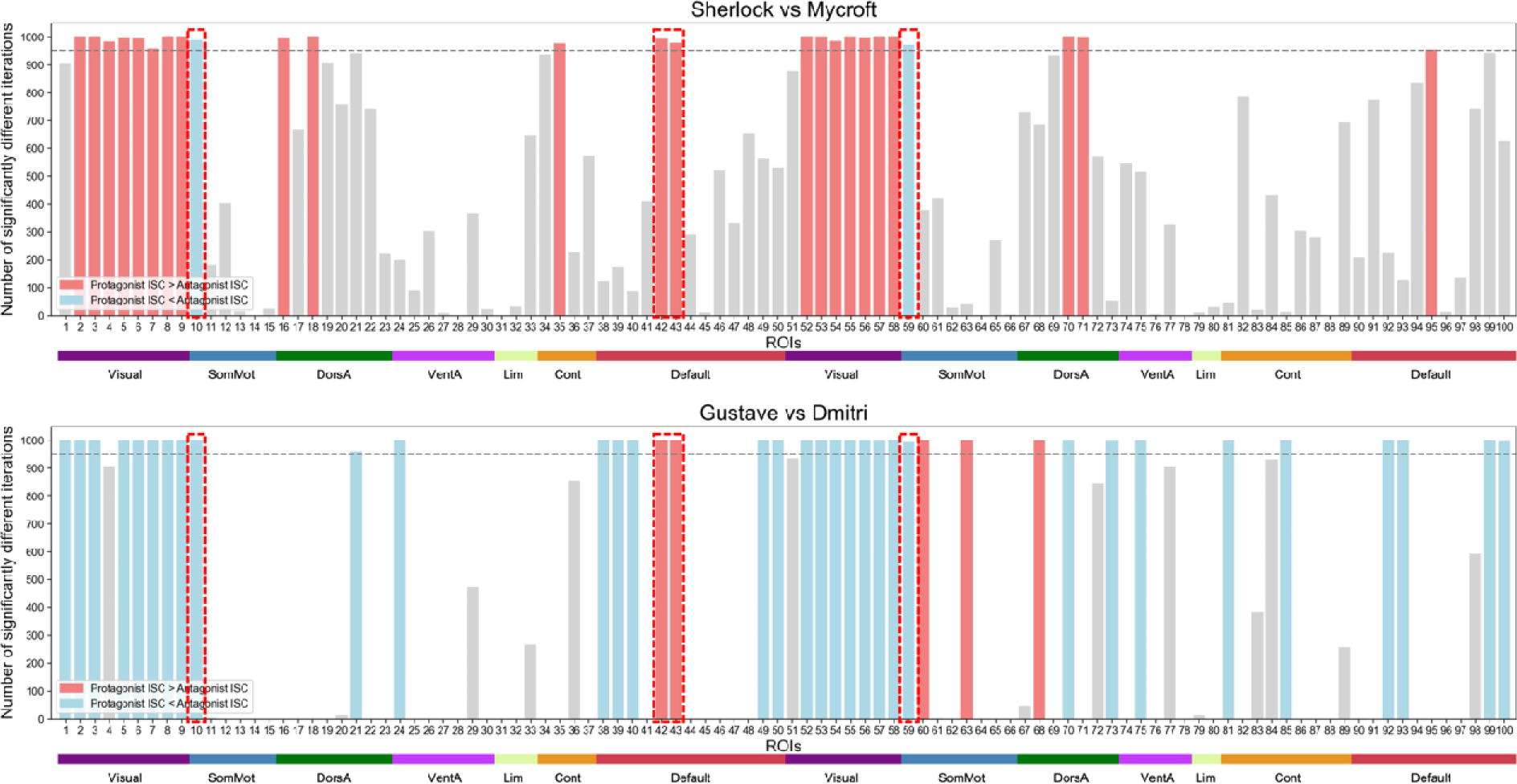
Summary of all ROIs with their number of iterations in which significant differences between protagonists and antagonists were observed. The horizontal gray dotted line marks 950 iterations. Gray bars represent the ROIs that the ISC values between the characters did not show consistent difference (> 95% of observations) from 1,000 iterations. Red bars represent ROIs whose ISC during the appearance of the protagonist was consistently greater than that of the antagonist, while blue bars show ROIs whose ISC during the appearance of the antagonist was consistently greater than that of the protagonist (> 95% of observations). The overlapping ROIs between the two datasets are highlighted in red boxes (parcels 10, 42, 43, and 59). The horizontal color bars show which of the Yeo networks the ROIs are categorized into. SomMot = Somatomotor, DorsA = Dorsal attention, VentA = Ventral attention, Lim = Limbic, Cont = Control.

**Figure 5.**
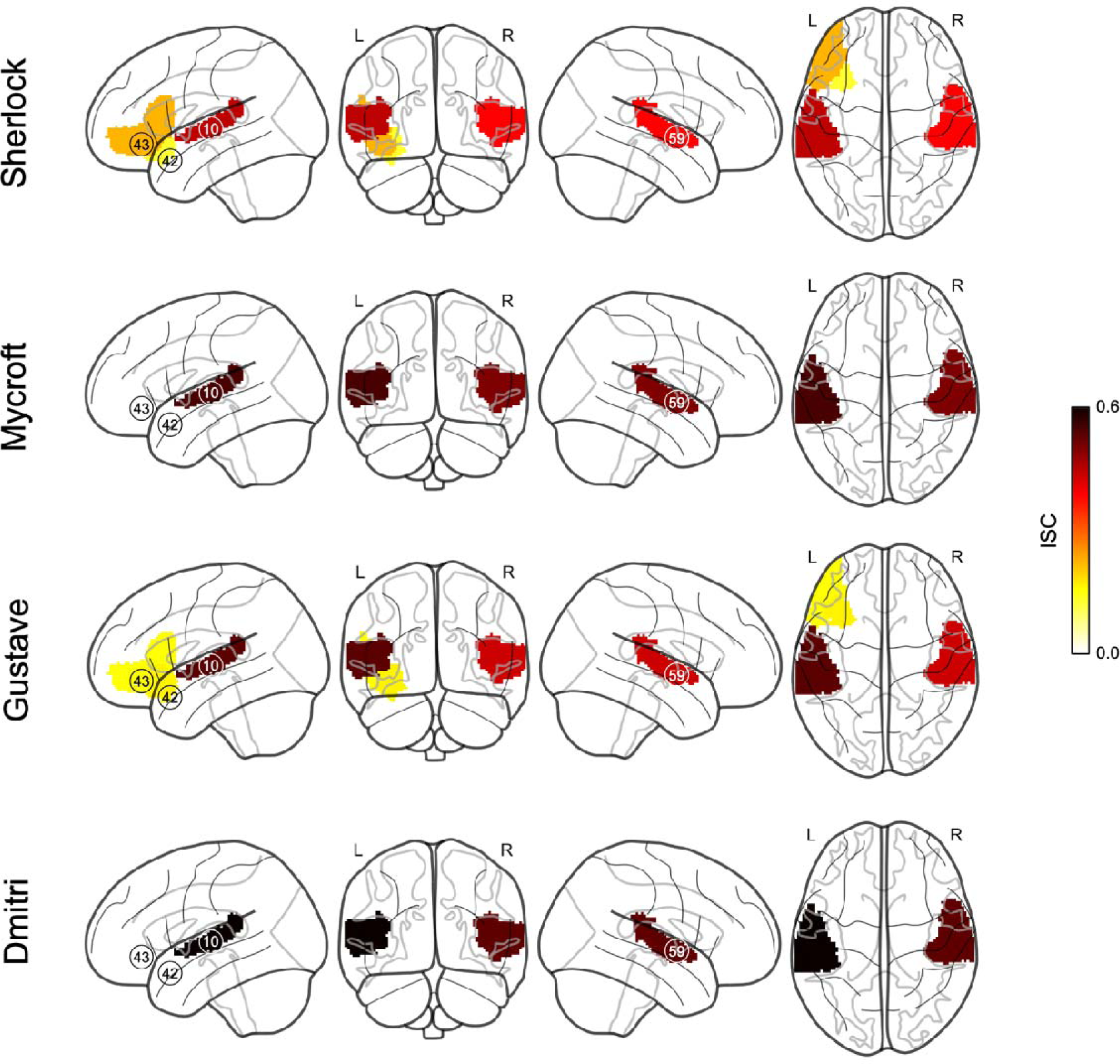
Parcels that showed distinctive synchronization pattern based on narrative roles (Bonferroni-corrected *p* < .0005). Darker colors represent greater ISC values. Across both datasets, parcels 42 and 43 showed greater synchronization in response to the protagonists (parcel 42: Sherlock ISC = 0.123, *p* = .0002; Mycroft ISC = 0.010, *ns*; Gustave ISC = 0.163, *p* = .0002; Dmitri ISC = 0.061, *ns*; parcel 43: Sherlock ISC = 0.217, *p* = .0002; Mycroft ISC = 0.072, *ns*; Gustave ISC = 0.145, *p* = .0002; Dmitri ISC = 0.036, *ns*), while parcels 10 and 59 showed greater synchronization in response to the antagonists (parcel 10: Sherlock ISC = 0.439, *p* = .0002; Mycroft ISC = 0.543, = .0002; Gustave ISC = 0.527, *p* = .0002; Dmitri ISC = 0.620, *p* = .0002; parcel 59: Sherlock ISC = 0.383, *p* = .0002; Mycroft ISC = 0.494, *p* = .0002; Gustave ISC = 0.426, *p* = .0002; Dmitri ISC = 0.522, *p* = .0002).

## Discussion

Here, we sought to identify brain regions that represent information about narrative roles by comparing protagonists and antagonists from two movie-watching functional neuroimaging datasets. Our analysis revealed increased neural synchronization in a network of brain regions including the inferior frontal gyrus (IFG) and orbitofrontal cortex (OFC) in response to protagonists. Meanwhile, activity of the bilateral auditory cortex and its neighboring regions showed higher inter-subject correlation (ISC) during the appearance of the antagonists. Notably, these findings were generalized across two independent datasets.

The three brain regions that showed greater neural synchronization in response to protagonists – IFG and OFC – are classified as part of the DMN (Schaefer et al., 2018). This puts our findings in proper context and offers a useful interpretation of the present findings. Traditionally, the DMN has been known to be activated when an individual is off-task, such as daydreaming or mind-wandering (Raichle et al., 2001; Fox et al., 2005). More recently, the DMN’s role has expanded to be involve narrative comprehension (Nguyen et al., 2019; Simony et al., 2016), self-referential thinking (Philippi et al., 2015; Ino et al., 2011), semantic information integration (Yang et al., 2023). and narrative engagement (Song et al., 2021; Ohad & Yeshurun, 2023). Our findings align particularly well with a recent fMRI study that showed individuals with similar DMN activity patterns with others were also the ones who engaged more with a positive character in a story (Ohad & Yeshurun, 2023). Our results further extend the findings from another fMRI study in which the DMN is suggested to support functions associated with social categorization by narrative roles (Ron et al., 2021). Specifically, we show that the DMN could be characterized by distinct synchronization patterns for the protagonists and antagonists, assuming that people generally tend to be more engaged towards the protagonists.

A key role of the IFG regarding narrative comprehension is semantic processing. The left IFG is associated with the level of immersion in a fictional narrative (Metz-Lutz et al., 2010), suggesting that entering the constructed reality in a play depends on verbal processing. The IFG is also related to the unification of conflicting prior world knowledge and given information (Hagoort et al., 2004; Yeshurun et al., 2017). Relatedly, based on the previous research showing that IFG activation increased in high-context conditions than in low-context conditions (Keidel et al., 2018), it can be inferred that the IFG may be involved in the integration of previous and new information (Thompson-Schill et al., 1997; Greenberg et al., 2005). Since the narrative revolves around the protagonists, the audience has more information about them compared to the antagonists. It follows then the increased neural synchronization in the IFG may reflect such information integration process, which occurs more frequently during the appearance of protagonists.

Furthermore, the IFG is a key node of the mirror neuron system, which has been suggested to be crucial in empathy, social behavior, and intention inference. Indeed, in line with the present findings, the IFG was involved while reading about protagonists and inferring their actions and intentions (Mason & Just, 2011). Interestingly, the activation of IFG is known to be more relevant in processing emotional empathy than cognitive empathy (Baird et al., 2010; Bodden et al., 2013; Schlaffke et al., 2015; Oliver et al., 2018; Pfeifer et al., 2008). Further supporting this notion, studies have found that individuals with lesions in the IFG area show poor performance in emotional empathy tasks (Shamay-Tsoory et al., 2009) or emotion inference tasks (Dal Monte et al., 2014). Assuming that people generally tend to show greater emotional empathy towards protagonists than antagonists in a story, we can speculate that our IFG results may be reflecting the differences in emotional empathy based on opposing narrative roles.

Along with the IFG, OFC is also involved in emotional perspective-taking (Hynes et al., 2006; Shamay-Tsoory et al., 2004) and emotional identification of voices or facial expressions (Hornak et al., 2003). Collectively, reduced volume in the left orbitofrontal and inferior frontal cortices in frontotemporal dementia was correlated with reduced empathic concern (Dermody et al., 2016). Thus, our results may be interpreted in this context, such that empathetic processing towards protagonists involves higher neural synchronization in regions such as the IFG and OFC.

On the other hand, regions that include mainly the bilateral auditory cortex (including Heschl’s gyrus, planum temporale, and parts of posterior and anterior superior temporal gyrus (STG)), the parietal operculum, and temporal pole showed higher synchronization during the appearance of the antagonists. The STG is well-known for its relation to auditory processing and speech perception (Chang et al., 2010; Mesgarani et al., 2014), but some studies have found it to support functions related to the perception of emotional content during audiovisual processing (Phillips et al., 1998), emotional learning (Grosso et al., 2015) and auditory memory (Weinberger, 2015). While it is difficult to pinpoint whether the bilateral parietal operculum and temporal pole were directly involved due to their close proximity to the auditory cortex, an interesting interpretation is possible considering the appearance of the antagonist typically entails threatening situations to the protagonist. The parietal operculum has been reported to exhibit increased activity when experiencing threatening stimuli (Straube & Miltner, 2011).

Furthermore, the parietal operculum is known to demonstrate distinct brain patterns based on the emotions elicited by external stimuli along with the STG (Sachs et al., 2018). In a similar vein, the temporal pole is often involved in selecting the most appropriate behavior in social situations. According to a review by Frith & Frith (2003), the temporal pole plays a role in generating a broader semantic and emotional context for the information being processed from past experiences. In social situations, this can be referred to as a ‘script’, which is a sequence of behavior that is considered proper and typical in a specific, well-known context (Schank & Abelson, 2013). As such, we could speculate that in a situation where the antagonists are posing a threat to the protagonists, the STG and its surrounding regions could elicit an emotional response and automatically infer the intention, computing the possible sequence of actions that the protagonists might take.

Consistent with previous studies, our findings elucidate that the brain regions within the DMN show differential synchronization patterns during the appearance of characters with different narrative roles. A few recent studies shed light on the neural representations of narrative roles (Ron et al., 2022; Ohad & Yeshurun, 2023) or fictional characters (Broom et al., 2021). However, these studies either utilized static visual stimuli or characters from a single narrative. To increase the generalizability of the findings and ecological validity, we used two independent datasets to identify and validate brain regions that show the same response based on narrative roles. As observed in the results from our analysis of the two datasets, neural synchronization patterns during movie-watching vary substantially depending on the characteristics of the movie. Future movie-watching fMRI studies would benefit from a validation step with multiple different narratives, especially when dealing with high-level social processes as in such cases the involvement of many brain regions would likely be dependent on the movie *per se*.

It is worth noting that we conducted an exploratory investigation to identify brain regions that exhibit differences in synchronization patterns based on the narrative roles, beyond the likability or valence of each character. Still, it is important to consider that some people might have a tendency to root for villains (Keen et al., 2021), and some narratives might have protagonists that are difficult to empathize with. While we were not able to test this possibility, future studies designed to take such factors into account would shed further light on the underlying mechanism for different narrative roles to engage different neural synchronization patterns.

One caveat of using a cortical parcellation atlas (Schaefer et al., 2018) is the omission of subcortical areas. Our additional analyses of the amygdala yielded no significant differences in synchronization, suggesting that unlike the IFG or OFC, the amygdala isn’t sensitive to information about narrative roles. In addition, inspecting physiological changes such as respiration, skin conductance, or heart rate variability could be helpful to understand the difference in emotional response to narrative roles in future studies (Saarimäki, 2021). With the limitation of using open datasets and a lack of behavioral data, we were not able to account for individual differences (Gruskin & Patel, 2022; Ohad & Yehshurun, 2023) such as tastes for each character or movie (Broom et al., 2021). Also, the present analysis was unable to address whether the dissociation in synchronization between the protagonists and antagonists might be due to emotional perception or emotional experience (Lindquist et al., 2012; Saarimäki, 2021).

In conclusion, we found that the IFG, and OFC, which are parts of the DMN, are conditionally involved in processing opposing narrative roles. A possible interpretation of these findings is that emotional empathy is automatically employed depending on a given character’s narrative roles during passive movie watching. Taken together, this study adds to the current literature that different narrative roles involve high-order social cognitive processing and may set the stage for a better understanding of person perception in more naturalistic settings.

## Acknowledgements

This research was supported by the National Research Foundation of Korea (NRF-2021R1F1A1045988). We thank Sujin Park and Chaebin Yoo for their helpful comments. We also thank the original authors of the *Sherlock* and *The Grand Budapest Hotel* datasets for their generosity in making it available for use.

## Author Contributions

H.R. and M.J.K. developed the study concept; H.R. analyzed the data under the supervision of M.J.K; H.R. and M.J.K. drafted the manuscript, and approved the final manuscript for submission.

