## supplementary for "Heroes and villains: opposing narrative roles engage neural synchronization in the lateral inferior frontal gyrus"

**Table S1.** Parcels with significant ISC (Bonferroni-corrected *p* < .0005) for Sherlock. Since ISC was calculated using subject-wise bootstrapping with 5,000 samples, the lowest possible *p*-value is .0002.

| **Parcel** | **Parcel label** | **ISC** | ***p*** |
| --- | --- | --- | --- |
| 1 | 7Networks_LH_Vis_1 | 0.242 | .0002 |
| 2 | 7Networks_LH_Vis_2 | 0.429 | .0002 |
| 3 | 7Networks_LH_Vis_3 | 0.495 | .0002 |
| 4 | 7Networks_LH_Vis_4 | 0.290 | .0002 |
| 5 | 7Networks_LH_Vis_5 | 0.334 | .0002 |
| 6 | 7Networks_LH_Vis_6 | 0.381 | .0002 |
| 7 | 7Networks_LH_Vis_7 | 0.487 | .0002 |
| 8 | 7Networks_LH_Vis_8 | 0.554 | .0002 |
| 9 | 7Networks_LH_Vis_9 | 0.344 | .0002 |
| 10 | 7Networks_LH_SomMot_1 | 0.439 | .0002 |
| 11 | 7Networks_LH_SomMot_2 | 0.092 | .0002 |
| 12 | 7Networks_LH_SomMot_3 | 0.113 | .0002 |
| 16 | 7Networks_LH_DorsAttn_Post_1 | 0.376 | .0002 |
| 17 | 7Networks_LH_DorsAttn_Post_2 | 0.177 | .0002 |
| 18 | 7Networks_LH_DorsAttn_Post_3 | 0.439 | .0002 |
| 19 | 7Networks_LH_DorsAttn_Post_4 | 0.194 | .0002 |
| 20 | 7Networks_LH_DorsAttn_Post_5 | 0.317 | .0002 |
| 21 | 7Networks_LH_DorsAttn_Post_6 | 0.249 | .0002 |
| 22 | 7Networks_LH_DorsAttn_PrCv_1 | 0.225 | .0002 |
| 23 | 7Networks_LH_DorsAttn_FEF_1 | 0.173 | .0002 |
| 24 | 7Networks_LH_SalVentAttn_ParOper_1 | 0.137 | .0002 |
| 25 | 7Networks_LH_SalVentAttn_FrOperIns_1 | 0.070 | .0002 |
| 26 | 7Networks_LH_SalVentAttn_FrOperIns_2 | 0.065 | .0002 |
| 27 | 7Networks_LH_SalVentAttn_PFCl_1 | 0.099 | .0002 |
| 28 | 7Networks_LH_SalVentAttn_Med_1 | 0.066 | .0002 |
| 29 | 7Networks_LH_SalVentAttn_Med_2 | 0.216 | .0002 |
| 30 | 7Networks_LH_SalVentAttn_Med_3 | 0.088 | .0002 |
| 31 | 7Networks_LH_Limbic_OFC_1 | 0.063 | .0002 |
| 32 | 7Networks_LH_Limbic_TempPole_1 | 0.091 | .0002 |
| 33 | 7Networks_LH_Limbic_TempPole_2 | 0.144 | .0002 |
| 34 | 7Networks_LH_Cont_Par_1 | 0.286 | .0002 |
| 35 | 7Networks_LH_Cont_PFCl_1 | 0.227 | .0002 |
| 36 | 7Networks_LH_Cont_pCun_1 | 0.291 | .0002 |
| 37 | 7Networks_LH_Cont_Cing_1 | 0.159 | .0002 |
| 38 | 7Networks_LH_Default_Temp_1 | 0.294 | .0002 |
| 39 | 7Networks_LH_Default_Temp_2 | 0.453 | .0002 |
| 40 | 7Networks_LH_Default_Par_1 | 0.313 | .0002 |
| 41 | 7Networks_LH_Default_Par_2 | 0.286 | .0002 |
| 42 | 7Networks_LH_Default_PFC_1 | 0.123 | .0002 |
| 43 | 7Networks_LH_Default_PFC_2 | 0.217 | .0002 |
| 44 | 7Networks_LH_Default_PFC_3 | 0.123 | .0002 |
| 46 | 7Networks_LH_Default_PFC_5 | 0.249 | .0002 |
| 47 | 7Networks_LH_Default_PFC_6 | 0.261 | .0002 |
| 48 | 7Networks_LH_Default_PFC_7 | 0.218 | .0002 |
| 49 | 7Networks_LH_Default_pCunPCC_1 | 0.251 | .0002 |
| 50 | 7Networks_LH_Default_pCunPCC_2 | 0.288 | .0002 |
| 51 | 7Networks_RH_Vis_1 | 0.240 | .0002 |
| 52 | 7Networks_RH_Vis_2 | 0.522 | .0002 |
| 53 | 7Networks_RH_Vis_3 | 0.367 | .0002 |
| 54 | 7Networks_RH_Vis_4 | 0.254 | .0002 |
| 55 | 7Networks_RH_Vis_5 | 0.465 | .0002 |
| 56 | 7Networks_RH_Vis_6 | 0.366 | .0002 |
| 57 | 7Networks_RH_Vis_7 | 0.587 | .0002 |
| 58 | 7Networks_RH_Vis_8 | 0.418 | .0002 |
| 59 | 7Networks_RH_SomMot_1 | 0.383 | .0002 |
| 60 | 7Networks_RH_SomMot_2 | 0.069 | .0002 |
| 61 | 7Networks_RH_SomMot_3 | 0.118 | .0002 |
| 63 | 7Networks_RH_SomMot_5 | 0.045 | .0002 |
| 65 | 7Networks_RH_SomMot_7 | 0.140 | .0002 |
| 66 | 7Networks_RH_SomMot_8 | 0.053 | .0002 |
| 67 | 7Networks_RH_DorsAttn_Post_1 | 0.311 | .0002 |
| 68 | 7Networks_RH_DorsAttn_Post_2 | 0.154 | .0002 |
| 69 | 7Networks_RH_DorsAttn_Post_3 | 0.294 | .0002 |
| 70 | 7Networks_RH_DorsAttn_Post_4 | 0.453 | .0002 |
| 71 | 7Networks_RH_DorsAttn_Post_5 | 0.304 | .0002 |
| 72 | 7Networks_RH_DorsAttn_PrCv_1 | 0.189 | .0002 |
| 73 | 7Networks_RH_DorsAttn_FEF_1 | 0.193 | .0002 |
| 74 | 7Networks_RH_SalVentAttn_TempOccPar_1 | 0.280 | .0002 |
| 75 | 7Networks_RH_SalVentAttn_TempOccPar_2 | 0.165 | .0002 |
| 76 | 7Networks_RH_SalVentAttn_FrOperIns_1 | 0.077 | .0002 |
| 77 | 7Networks_RH_SalVentAttn_Med_1 | 0.179 | .0002 |
| 78 | 7Networks_RH_SalVentAttn_Med_2 | 0.078 | .0002 |
| 79 | 7Networks_RH_Limbic_OFC_1 | 0.067 | .0002 |
| 80 | 7Networks_RH_Limbic_TempPole_1 | 0.076 | .0002 |
| 81 | 7Networks_RH_Cont_Par_1 | 0.170 | .0002 |
| 82 | 7Networks_RH_Cont_Par_2 | 0.260 | .0002 |
| 84 | 7Networks_RH_Cont_PFCl_2 | 0.190 | .0002 |
| 85 | 7Networks_RH_Cont_PFCl_3 | 0.106 | .0002 |
| 86 | 7Networks_RH_Cont_PFCl_4 | 0.202 | .0002 |
| 87 | 7Networks_RH_Cont_Cing_1 | 0.152 | .0002 |
| 88 | 7Networks_RH_Cont_PFCmp_1 | 0.082 | .0002 |
| 89 | 7Networks_RH_Cont_pCun_1 | 0.338 | .0002 |
| 90 | 7Networks_RH_Default_Par_1 | 0.248 | .0002 |
| 91 | 7Networks_RH_Default_Temp_1 | 0.147 | .0002 |
| 92 | 7Networks_RH_Default_Temp_2 | 0.212 | .0002 |
| 93 | 7Networks_RH_Default_Temp_3 | 0.385 | .0002 |
| 94 | 7Networks_RH_Default_PFCv_1 | 0.091 | .0002 |
| 95 | 7Networks_RH_Default_PFCv_2 | 0.163 | .0002 |
| 96 | 7Networks_RH_Default_PFCdPFCm_1 | 0.126 | .0002 |
| 97 | 7Networks_RH_Default_PFCdPFCm_2 | 0.198 | .0002 |
| 98 | 7Networks_RH_Default_PFCdPFCm_3 | 0.214 | .0002 |
| 99 | 7Networks_RH_Default_pCunPCC_1 | 0.333 | .0002 |
| 100 | 7Networks_RH_Default_pCunPCC_2 | 0.294 | .0002 |

**Table S2.** Parcels with significant ISC (Bonferroni-corrected *p* < .0005) for Mycroft.

| **Parcel** | **Parcel label** | **ISC** | ***p*** |
| --- | --- | --- | --- |
| 1 | 7Networks_LH_Vis_1 | 0.134 | .0002 |
| 2 | 7Networks_LH_Vis_2 | 0.165 | .0002 |
| 3 | 7Networks_LH_Vis_3 | 0.258 | .0002 |
| 4 | 7Networks_LH_Vis_4 | 0.124 | .0002 |
| 5 | 7Networks_LH_Vis_5 | 0.157 | .0002 |
| 6 | 7Networks_LH_Vis_6 | 0.233 | .0002 |
| 7 | 7Networks_LH_Vis_7 | 0.386 | .0002 |
| 8 | 7Networks_LH_Vis_8 | 0.290 | .0002 |
| 10 | 7Networks_LH_SomMot_1 | 0.543 | .0002 |
| 11 | 7Networks_LH_SomMot_2 | 0.138 | .0002 |
| 12 | 7Networks_LH_SomMot_3 | 0.167 | .0002 |
| 16 | 7Networks_LH_DorsAttn_Post_1 | 0.185 | .0002 |
| 20 | 7Networks_LH_DorsAttn_Post_5 | 0.249 | .0002 |
| 21 | 7Networks_LH_DorsAttn_Post_6 | 0.117 | .0002 |
| 24 | 7Networks_LH_SalVentAttn_ParOper_1 | 0.121 | .0002 |
| 29 | 7Networks_LH_SalVentAttn_Med_2 | 0.148 | .0002 |
| 33 | 7Networks_LH_Limbic_TempPole_2 | 0.077 | .0002 |
| 34 | 7Networks_LH_Cont_Par_1 | 0.173 | .0002 |
| 36 | 7Networks_LH_Cont_pCun_1 | 0.303 | .0002 |
| 38 | 7Networks_LH_Default_Temp_1 | 0.289 | .0002 |
| 39 | 7Networks_LH_Default_Temp_2 | 0.447 | .0002 |
| 40 | 7Networks_LH_Default_Par_1 | 0.269 | .0002 |
| 41 | 7Networks_LH_Default_Par_2 | 0.226 | .0002 |
| 46 | 7Networks_LH_Default_PFC_5 | 0.176 | .0002 |
| 47 | 7Networks_LH_Default_PFC_6 | 0.207 | .0002 |
| 48 | 7Networks_LH_Default_PFC_7 | 0.147 | .0002 |
| 49 | 7Networks_LH_Default_pCunPCC_1 | 0.192 | .0002 |
| 50 | 7Networks_LH_Default_pCunPCC_2 | 0.249 | .0002 |
| 51 | 7Networks_RH_Vis_1 | 0.146 | .0002 |
| 52 | 7Networks_RH_Vis_2 | 0.296 | .0002 |
| 53 | 7Networks_RH_Vis_3 | 0.210 | .0002 |
| 54 | 7Networks_RH_Vis_4 | 0.103 | .0002 |
| 55 | 7Networks_RH_Vis_5 | 0.280 | .0002 |
| 56 | 7Networks_RH_Vis_6 | 0.225 | .0002 |
| 57 | 7Networks_RH_Vis_7 | 0.337 | .0002 |
| 58 | 7Networks_RH_Vis_8 | 0.198 | .0002 |
| 59 | 7Networks_RH_SomMot_1 | 0.494 | .0002 |
| 61 | 7Networks_RH_SomMot_3 | 0.176 | .0002 |
| 67 | 7Networks_RH_DorsAttn_Post_1 | 0.229 | .0002 |
| 69 | 7Networks_RH_DorsAttn_Post_3 | 0.152 | .0002 |
| 70 | 7Networks_RH_DorsAttn_Post_4 | 0.208 | .0002 |
| 71 | 7Networks_RH_DorsAttn_Post_5 | 0.122 | .0002 |
| 73 | 7Networks_RH_DorsAttn_FEF_1 | 0.158 | .0002 |
| 74 | 7Networks_RH_SalVentAttn_TempOccPar_1 | 0.214 | .0002 |
| 75 | 7Networks_RH_SalVentAttn_TempOccPar_2 | 0.116 | .0002 |
| 76 | 7Networks_RH_SalVentAttn_FrOperIns_1 | 0.113 | .0002 |
| 81 | 7Networks_RH_Cont_Par_1 | 0.170 | .0002 |
| 82 | 7Networks_RH_Cont_Par_2 | 0.144 | .0002 |
| 85 | 7Networks_RH_Cont_PFCl_3 | 0.148 | .0002 |
| 88 | 7Networks_RH_Cont_PFCmp_1 | 0.086 | .0002 |
| 89 | 7Networks_RH_Cont_pCun_1 | 0.281 | .0002 |
| 90 | 7Networks_RH_Default_Par_1 | 0.181 | .0002 |
| 92 | 7Networks_RH_Default_Temp_2 | 0.177 | .0002 |
| 93 | 7Networks_RH_Default_Temp_3 | 0.375 | .0002 |
| 96 | 7Networks_RH_Default_PFCdPFCm_1 | 0.128 | .0002 |
| 97 | 7Networks_RH_Default_PFCdPFCm_2 | 0.152 | .0002 |
| 98 | 7Networks_RH_Default_PFCdPFCm_3 | 0.105 | .0002 |
| 99 | 7Networks_RH_Default_pCunPCC_1 | 0.196 | .0002 |
| 100 | 7Networks_RH_Default_pCunPCC_2 | 0.224 | .0002 |

**Table S3.** Parcels with significant ISC (Bonferroni-corrected *p* < .0005) for Gustave.

| **Parcel** | **Parcel label** | **ISC** | ***p*** |
| --- | --- | --- | --- |
| 1 | 7Networks_LH_Vis_1 | 0.197 | .0002 |
| 2 | 7Networks_LH_Vis_2 | 0.329 | .0002 |
| 3 | 7Networks_LH_Vis_3 | 0.164 | .0002 |
| 4 | 7Networks_LH_Vis_4 | 0.258 | .0002 |
| 5 | 7Networks_LH_Vis_5 | 0.217 | .0002 |
| 6 | 7Networks_LH_Vis_6 | 0.197 | .0002 |
| 7 | 7Networks_LH_Vis_7 | 0.385 | .0002 |
| 8 | 7Networks_LH_Vis_8 | 0.330 | .0002 |
| 9 | 7Networks_LH_Vis_9 | 0.112 | .0002 |
| 10 | 7Networks_LH_SomMot_1 | 0.527 | .0002 |
| 11 | 7Networks_LH_SomMot_2 | 0.187 | .0002 |
| 12 | 7Networks_LH_SomMot_3 | 0.172 | .0002 |
| 13 | 7Networks_LH_SomMot_4 | 0.061 | .0002 |
| 15 | 7Networks_LH_SomMot_6 | 0.063 | .0002 |
| 16 | 7Networks_LH_DorsAttn_Post_1 | 0.240 | .0002 |
| 17 | 7Networks_LH_DorsAttn_Post_2 | 0.144 | .0002 |
| 18 | 7Networks_LH_DorsAttn_Post_3 | 0.283 | .0002 |
| 19 | 7Networks_LH_DorsAttn_Post_4 | 0.119 | .0002 |
| 20 | 7Networks_LH_DorsAttn_Post_5 | 0.248 | .0002 |
| 21 | 7Networks_LH_DorsAttn_Post_6 | 0.154 | .0002 |
| 22 | 7Networks_LH_DorsAttn_PrCv_1 | 0.073 | .0002 |
| 23 | 7Networks_LH_DorsAttn_FEF_1 | 0.118 | .0002 |
| 25 | 7Networks_LH_SalVentAttn_FrOperIns_1 | 0.066 | .0002 |
| 26 | 7Networks_LH_SalVentAttn_FrOperIns_2 | 0.065 | .0002 |
| 28 | 7Networks_LH_SalVentAttn_Med_1 | 0.091 | .0002 |
| 29 | 7Networks_LH_SalVentAttn_Med_2 | 0.124 | .0002 |
| 34 | 7Networks_LH_Cont_Par_1 | 0.150 | .0002 |
| 36 | 7Networks_LH_Cont_pCun_1 | 0.101 | .0002 |
| 37 | 7Networks_LH_Cont_Cing_1 | 0.089 | .0002 |
| 38 | 7Networks_LH_Default_Temp_1 | 0.369 | .0002 |
| 39 | 7Networks_LH_Default_Temp_2 | 0.447 | .0002 |
| 40 | 7Networks_LH_Default_Par_1 | 0.310 | .0002 |
| 41 | 7Networks_LH_Default_Par_2 | 0.172 | .0002 |
| 42 | 7Networks_LH_Default_PFC_1 | 0.163 | .0002 |
| 43 | 7Networks_LH_Default_PFC_2 | 0.145 | .0002 |
| 45 | 7Networks_LH_Default_PFC_4 | 0.112 | .0002 |
| 46 | 7Networks_LH_Default_PFC_5 | 0.159 | .0002 |
| 47 | 7Networks_LH_Default_PFC_6 | 0.133 | .0002 |
| 48 | 7Networks_LH_Default_PFC_7 | 0.081 | .0002 |
| 50 | 7Networks_LH_Default_pCunPCC_2 | 0.168 | .0002 |
| 51 | 7Networks_RH_Vis_1 | 0.258 | .0002 |
| 52 | 7Networks_RH_Vis_2 | 0.350 | .0002 |
| 53 | 7Networks_RH_Vis_3 | 0.346 | .0002 |
| 54 | 7Networks_RH_Vis_4 | 0.175 | .0002 |
| 55 | 7Networks_RH_Vis_5 | 0.196 | .0002 |
| 56 | 7Networks_RH_Vis_6 | 0.115 | .0002 |
| 57 | 7Networks_RH_Vis_7 | 0.397 | .0002 |
| 58 | 7Networks_RH_Vis_8 | 0.144 | .0002 |
| 59 | 7Networks_RH_SomMot_1 | 0.426 | .0002 |
| 60 | 7Networks_RH_SomMot_2 | 0.136 | .0002 |
| 61 | 7Networks_RH_SomMot_3 | 0.185 | .0002 |
| 63 | 7Networks_RH_SomMot_5 | 0.116 | .0002 |
| 65 | 7Networks_RH_SomMot_7 | 0.120 | .0002 |
| 66 | 7Networks_RH_SomMot_8 | 0.058 | .0002 |
| 67 | 7Networks_RH_DorsAttn_Post_1 | 0.359 | .0002 |
| 68 | 7Networks_RH_DorsAttn_Post_2 | 0.261 | .0002 |
| 69 | 7Networks_RH_DorsAttn_Post_3 | 0.210 | .0002 |
| 70 | 7Networks_RH_DorsAttn_Post_4 | 0.244 | .0002 |
| 71 | 7Networks_RH_DorsAttn_Post_5 | 0.239 | .0002 |
| 72 | 7Networks_RH_DorsAttn_PrCv_1 | 0.123 | .0002 |
| 73 | 7Networks_RH_DorsAttn_FEF_1 | 0.159 | .0002 |
| 74 | 7Networks_RH_SalVentAttn_TempOccPar_1 | 0.357 | .0002 |
| 77 | 7Networks_RH_SalVentAttn_Med_1 | 0.128 | .0002 |
| 80 | 7Networks_RH_Limbic_TempPole_1 | 0.093 | .0002 |
| 81 | 7Networks_RH_Cont_Par_1 | 0.162 | .0002 |
| 82 | 7Networks_RH_Cont_Par_2 | 0.172 | .0002 |
| 83 | 7Networks_RH_Cont_PFCl_1 | 0.127 | .0002 |
| 85 | 7Networks_RH_Cont_PFCl_3 | 0.080 | .0002 |
| 86 | 7Networks_RH_Cont_PFCl_4 | 0.148 | .0002 |
| 89 | 7Networks_RH_Cont_pCun_1 | 0.191 | .0002 |
| 90 | 7Networks_RH_Default_Par_1 | 0.284 | .0002 |
| 91 | 7Networks_RH_Default_Temp_1 | 0.198 | .0002 |
| 92 | 7Networks_RH_Default_Temp_2 | 0.316 | .0002 |
| 93 | 7Networks_RH_Default_Temp_3 | 0.494 | .0002 |
| 94 | 7Networks_RH_Default_PFCv_1 | 0.139 | .0002 |
| 95 | 7Networks_RH_Default_PFCv_2 | 0.188 | .0002 |
| 97 | 7Networks_RH_Default_PFCdPFCm_2 | 0.225 | .0002 |
| 98 | 7Networks_RH_Default_PFCdPFCm_3 | 0.109 | .0002 |
| 99 | 7Networks_RH_Default_pCunPCC_1 | 0.131 | .0002 |
| 100 | 7Networks_RH_Default_pCunPCC_2 | 0.180 | .0002 |

**Table S4.** Parcels with significant ISC (Bonferroni-corrected *p* < .0005) for Dmitri.

| **Parcel** | **Parcel label** | **ISC** | ***p*** |
| --- | --- | --- | --- |
| 1 | 7Networks_LH_Vis_1 | 0.302 | .0002 |
| 2 | 7Networks_LH_Vis_2 | 0.457 | .0002 |
| 3 | 7Networks_LH_Vis_3 | 0.388 | .0002 |
| 4 | 7Networks_LH_Vis_4 | 0.318 | .0002 |
| 5 | 7Networks_LH_Vis_5 | 0.396 | .0002 |
| 6 | 7Networks_LH_Vis_6 | 0.327 | .0002 |
| 7 | 7Networks_LH_Vis_7 | 0.326 | .0002 |
| 8 | 7Networks_LH_Vis_8 | 0.467 | .0002 |
| 9 | 7Networks_LH_Vis_9 | 0.240 | .0002 |
| 10 | 7Networks_LH_SomMot_1 | 0.620 | .0002 |
| 11 | 7Networks_LH_SomMot_2 | 0.125 | .0002 |
| 12 | 7Networks_LH_SomMot_3 | 0.190 | .0002 |
| 16 | 7Networks_LH_DorsAttn_Post_1 | 0.213 | .0002 |
| 17 | 7Networks_LH_DorsAttn_Post_2 | 0.182 | .0002 |
| 18 | 7Networks_LH_DorsAttn_Post_3 | 0.264 | .0002 |
| 19 | 7Networks_LH_DorsAttn_Post_4 | 0.119 | .0002 |
| 20 | 7Networks_LH_DorsAttn_Post_5 | 0.220 | .0002 |
| 21 | 7Networks_LH_DorsAttn_Post_6 | 0.252 | .0002 |
| 23 | 7Networks_LH_DorsAttn_FEF_1 | 0.140 | .0002 |
| 24 | 7Networks_LH_SalVentAttn_ParOper_1 | 0.177 | .0002 |
| 25 | 7Networks_LH_SalVentAttn_FrOperIns_1 | 0.075 | .0002 |
| 27 | 7Networks_LH_SalVentAttn_PFCl_1 | 0.084 | .0002 |
| 28 | 7Networks_LH_SalVentAttn_Med_1 | 0.077 | .0002 |
| 29 | 7Networks_LH_SalVentAttn_Med_2 | 0.173 | .0002 |
| 30 | 7Networks_LH_SalVentAttn_Med_3 | 0.078 | .0002 |
| 32 | 7Networks_LH_Limbic_TempPole_1 | 0.090 | .0002 |
| 34 | 7Networks_LH_Cont_Par_1 | 0.126 | .0002 |
| 36 | 7Networks_LH_Cont_pCun_1 | 0.151 | .0002 |
| 38 | 7Networks_LH_Default_Temp_1 | 0.491 | .0002 |
| 39 | 7Networks_LH_Default_Temp_2 | 0.594 | .0002 |
| 40 | 7Networks_LH_Default_Par_1 | 0.486 | .0002 |
| 41 | 7Networks_LH_Default_Par_2 | 0.210 | .0002 |
| 44 | 7Networks_LH_Default_PFC_3 | 0.069 | .0002 |
| 45 | 7Networks_LH_Default_PFC_4 | 0.116 | .0002 |
| 46 | 7Networks_LH_Default_PFC_5 | 0.157 | .0002 |
| 48 | 7Networks_LH_Default_PFC_7 | 0.116 | .0002 |
| 49 | 7Networks_LH_Default_pCunPCC_1 | 0.199 | .0002 |
| 50 | 7Networks_LH_Default_pCunPCC_2 | 0.245 | .0002 |
| 51 | 7Networks_RH_Vis_1 | 0.296 | .0002 |
| 52 | 7Networks_RH_Vis_2 | 0.522 | .0002 |
| 53 | 7Networks_RH_Vis_3 | 0.276 | .0002 |
| 54 | 7Networks_RH_Vis_4 | 0.369 | .0002 |
| 55 | 7Networks_RH_Vis_5 | 0.418 | .0002 |
| 56 | 7Networks_RH_Vis_6 | 0.289 | .0002 |
| 57 | 7Networks_RH_Vis_7 | 0.491 | .0002 |
| 58 | 7Networks_RH_Vis_8 | 0.339 | .0002 |
| 59 | 7Networks_RH_SomMot_1 | 0.522 | .0002 |
| 60 | 7Networks_RH_SomMot_2 | 0.095 | .0002 |
| 61 | 7Networks_RH_SomMot_3 | 0.155 | .0002 |
| 65 | 7Networks_RH_SomMot_7 | 0.144 | .0002 |
| 67 | 7Networks_RH_DorsAttn_Post_1 | 0.313 | .0002 |
| 68 | 7Networks_RH_DorsAttn_Post_2 | 0.150 | .0002 |
| 69 | 7Networks_RH_DorsAttn_Post_3 | 0.187 | .0002 |
| 70 | 7Networks_RH_DorsAttn_Post_4 | 0.331 | .0002 |
| 71 | 7Networks_RH_DorsAttn_Post_5 | 0.270 | .0002 |
| 72 | 7Networks_RH_DorsAttn_PrCv_1 | 0.166 | .0002 |
| 73 | 7Networks_RH_DorsAttn_FEF_1 | 0.264 | .0002 |
| 74 | 7Networks_RH_SalVentAttn_TempOccPar_1 | 0.359 | .0002 |
| 75 | 7Networks_RH_SalVentAttn_TempOccPar_2 | 0.205 | .0002 |
| 77 | 7Networks_RH_SalVentAttn_Med_1 | 0.177 | .0002 |
| 78 | 7Networks_RH_SalVentAttn_Med_2 | 0.091 | .0002 |
| 80 | 7Networks_RH_Limbic_TempPole_1 | 0.094 | .0002 |
| 81 | 7Networks_RH_Cont_Par_1 | 0.267 | .0002 |
| 82 | 7Networks_RH_Cont_Par_2 | 0.206 | .0002 |
| 83 | 7Networks_RH_Cont_PFCl_1 | 0.175 | .0002 |
| 84 | 7Networks_RH_Cont_PFCl_2 | 0.136 | .0002 |
| 85 | 7Networks_RH_Cont_PFCl_3 | 0.165 | .0002 |
| 86 | 7Networks_RH_Cont_PFCl_4 | 0.194 | .0002 |
| 87 | 7Networks_RH_Cont_Cing_1 | 0.119 | .0002 |
| 88 | 7Networks_RH_Cont_PFCmp_1 | 0.101 | .0002 |
| 89 | 7Networks_RH_Cont_pCun_1 | 0.252 | .0002 |
| 90 | 7Networks_RH_Default_Par_1 | 0.249 | .0002 |
| 91 | 7Networks_RH_Default_Temp_1 | 0.216 | .0002 |
| 92 | 7Networks_RH_Default_Temp_2 | 0.448 | .0002 |
| 93 | 7Networks_RH_Default_Temp_3 | 0.625 | .0002 |
| 94 | 7Networks_RH_Default_PFCv_1 | 0.106 | .0002 |
| 95 | 7Networks_RH_Default_PFCv_2 | 0.153 | .0002 |
| 96 | 7Networks_RH_Default_PFCdPFCm_1 | 0.110 | .0002 |
| 97 | 7Networks_RH_Default_PFCdPFCm_2 | 0.215 | .0002 |
| 98 | 7Networks_RH_Default_PFCdPFCm_3 | 0.176 | .0002 |
| 99 | 7Networks_RH_Default_pCunPCC_1 | 0.302 | .0002 |
| 100 | 7Networks_RH_Default_pCunPCC_2 | 0.457 | .0002 |

**Table S5.** Parcels with significant ISC differences (Bonferroni-corrected *p* < .0005) between the protagonist vs. antagonist (i.e., Sherlock and Mycroft) in *Sherlock*. Paired *t*-tests were repeated 1,000 times with resampled TRs, and the parcels that differed significantly for more than 950 iterations are summarized below.

| **Parcel** | **Parcel label** | **Sherlock** | | **Mycroft** | |
| --- | --- | --- | --- | --- | --- |
|  |  | **ISC** | ***p*** | **ISC** | ***p*** |
| 2 | 7Networks_LH_Vis_2 | 0.429 | .0002 | 0.165 | .0002 |
| 3 | 7Networks_LH_Vis_3 | 0.495 | .0002 | 0.258 | .0002 |
| 4 | 7Networks_LH_Vis_4 | 0.290 | .0002 | 0.124 | .0002 |
| 5 | 7Networks_LH_Vis_5 | 0.334 | .0002 | 0.157 | .0002 |
| 6 | 7Networks_LH_Vis_6 | 0.381 | .0002 | 0.233 | .0002 |
| 7 | 7Networks_LH_Vis_7 | 0.487 | .0002 | 0.386 | .0002 |
| 8 | 7Networks_LH_Vis_8 | 0.554 | .0002 | 0.290 | .0002 |
| 9 | 7Networks_LH_Vis_9 | 0.344 | .0002 | 0.169 | .0002 |
| 10 | 7Networks_LH_SomMot_1 | 0.439 | .0002 | 0.543 | .0002 |
| 16 | 7Networks_LH_DorsAttn_Post_1 | 0.376 | .0002 | 0.185 | .0002 |
| 18 | 7Networks_LH_DorsAttn_Post_3 | 0.439 | .0002 | 0.233 | .0002 |
| 35 | 7Networks_LH_Cont_PFCl_1 | 0.227 | .0002 | 0.074 | *ns* |
| 42 | 7Networks_LH_Default_PFC_1 | 0.123 | .0002 | 0.010 | *ns* |
| 43 | 7Networks_LH_Default_PFC_2 | 0.217 | .0002 | 0.072 | *ns* |
| 52 | 7Networks_RH_Vis_2 | 0.521 | .0002 | 0.296 | .0002 |
| 53 | 7Networks_RH_Vis_3 | 0.367 | .0002 | 0.210 | .0002 |
| 54 | 7Networks_RH_Vis_4 | 0.254 | .0002 | 0.103 | .0002 |
| 55 | 7Networks_RH_Vis_5 | 0.465 | .0002 | 0.280 | .0002 |
| 56 | 7Networks_RH_Vis_6 | 0.366 | .0002 | 0.225 | .0002 |
| 57 | 7Networks_RH_Vis_7 | 0.587 | .0002 | 0.337 | .0002 |
| 58 | 7Networks_RH_Vis_8 | 0.418 | .0002 | 0.198 | .0002 |
| 59 | 7Networks_RH_SomMot_1 | 0.383 | .0002 | 0.494 | .0002 |
| 70 | 7Networks_RH_DorsAttn_Post_4 | 0.453 | .0002 | 0.208 | .0002 |
| 71 | 7Networks_RH_DorsAttn_Post_5 | 0.304 | .0002 | 0.122 | *ns* |
| 95 | 7Networks_RH_Default_PFCv_2 | 0.163 | .0002 | 0.061 | *ns* |

**Table S6.** Parcels with significant ISC differences (Bonferroni-corrected *p* < .0005) between the protagonist vs. antagonist (i.e., Gustave and Dmitri) in *The Grand Budapest Hotel*. Paired *t*-tests were repeated 1,000 times with resampled TRs, and the parcels that differed significantly for more than 950 iterations are summarized below.

| **Parcel** | **Parcel label** | **Gustave** | | **Dmitri** | |
| --- | --- | --- | --- | --- | --- |
|  |  | **ISC** | ***p*** | **ISC** | ***p*** |
| 1 | 7Networks_LH_Vis_1 | 0.197 | .0002 | 0.302 | .0002 |
| 2 | 7Networks_LH_Vis_2 | 0.329 | .0002 | 0.457 | .0002 |
| 3 | 7Networks_LH_Vis_3 | 0.164 | .0002 | 0.388 | .0002 |
| 5 | 7Networks_LH_Vis_5 | 0.217 | .0002 | 0.396 | .0002 |
| 6 | 7Networks_LH_Vis_6 | 0.197 | .0002 | 0.327 | .0002 |
| 7 | 7Networks_LH_Vis_7 | 0.385 | .0002 | 0.326 | .0002 |
| 8 | 7Networks_LH_Vis_8 | 0.330 | .0002 | 0.467 | .0002 |
| 9 | 7Networks_LH_Vis_9 | 0.112 | .0002 | 0.240 | .0002 |
| 10 | 7Networks_LH_SomMot_1 | 0.527 | .0002 | 0.620 | .0002 |
| 21 | 7Networks_LH_DorsAttn_Post_6 | 0.154 | .0002 | 0.252 | .0002 |
| 24 | 7Networks_LH_SalVentAttn_ParOper_1 | 0.046 | *ns* | 0.177 | .0002 |
| 38 | 7Networks_LH_Default_Temp_1 | 0.369 | .0002 | 0.491 | .0002 |
| 39 | 7Networks_LH_Default_Temp_2 | 0.447 | .0002 | 0.594 | .0002 |
| 40 | 7Networks_LH_Default_Par_1 | 0.310 | .0002 | 0.486 | .0002 |
| 42 | 7Networks_LH_Default_PFC_1 | 0.163 | .0002 | 0.061 | *ns* |
| 43 | 7Networks_LH_Default_PFC_2 | 0.145 | .0002 | 0.036 | *ns* |
| 49 | 7Networks_LH_Default_pCunPCC_1 | 0.042 | *ns* | 0.199 | .0002 |
| 50 | 7Networks_LH_Default_pCunPCC_2 | 0.168 | .0002 | 0.245 | .0002 |
| 52 | 7Networks_RH_Vis_2 | 0.350 | .0002 | 0.522 | .0002 |
| 53 | 7Networks_RH_Vis_3 | 0.346 | .0002 | 0.276 | .0002 |
| 54 | 7Networks_RH_Vis_4 | 0.175 | .0002 | 0.369 | .0002 |
| 55 | 7Networks_RH_Vis_5 | 0.196 | .0002 | 0.418 | .0002 |
| 56 | 7Networks_RH_Vis_6 | 0.115 | .0002 | 0.289 | .0002 |
| 57 | 7Networks_RH_Vis_7 | 0.397 | .0002 | 0.491 | .0002 |
| 58 | 7Networks_RH_Vis_8 | 0.144 | .0002 | 0.339 | .0002 |
| 59 | 7Networks_RH_SomMot_1 | 0.426 | .0002 | 0.522 | .0002 |
| 60 | 7Networks_RH_SomMot_2 | 0.136 | .0002 | 0.095 | .0002 |
| 63 | 7Networks_RH_SomMot_5 | 0.116 | .0002 | 0.045 | *ns* |
| 68 | 7Networks_RH_DorsAttn_Post_2 | 0.261 | .0002 | 0.150 | .0002 |
| 70 | 7Networks_RH_DorsAttn_Post_4 | 0.244 | .0002 | 0.331 | .0002 |
| 73 | 7Networks_RH_DorsAttn_FEF_1 | 0.159 | .0002 | 0.264 | .0002 |
| 75 | 7Networks_RH_SalVentAttn_TempOccPar_2 | 0.074 | *ns* | 0.205 | .0002 |
| 81 | 7Networks_RH_Cont_Par_1 | 0.162 | .0002 | 0.267 | .0002 |
| 85 | 7Networks_RH_Cont_PFCl_3 | 0.080 | .0002 | 0.165 | .0002 |
| 92 | 7Networks_RH_Default_Temp_2 | 0.316 | .0002 | 0.448 | .0002 |
| 93 | 7Networks_RH_Default_Temp_3 | 0.494 | .0002 | 0.625 | .0002 |
| 99 | 7Networks_RH_Default_pCunPCC_1 | 0.130 | .0002 | 0.258 | .0002 |
| 100 | 7Networks_RH_Default_pCunPCC_2 | 0.180 | .0002 | 0.253 | .0002 |
